# Isocaloric restriction normalizes body weight but not relative adiposity, adipose inflammation, or gut microbial diversity in high-fat diet-fed mice

**DOI:** 10.64898/2026.10.23.761336

**Authors:** Sungjin Eun, Eunji Jeong, Seoyeon Chae, Sunhye Lee

## Abstract

**Background/Objectives:** The separate contributions of excess energy intake and diet composition to high-fat diet (HFD)-induced adiposity, adipose inflammation, and gut microbial changes are unclear. We tested whether restricting HFD intake to the energy intake of low-fat (LF)-fed mice (isocaloric restriction) prevents or reverses these outcomes.

**Subjects/Methods:** Male C57BL/6 mice (n = 8/group) received for 12 weeks an LF diet (LF/LF) or HFD (HF/HF) ad libitum, HFD ad libitum then isocaloric HFD from week 7 (HF/IHF), or isocaloric HFD throughout (IHF/IHF). Adiposity, plasma markers, adipose and ileal gene expression, and fecal microbiota (16S rRNA sequencing, n = 4/group) were assessed.

**Results:** Final body weight in IHF/IHF did not differ from LF/LF (difference 0.1 g, 95% CI −6.5 to 6.7; *P* > 0.99) and was lower than in HF/HF (−8.2 g, −14.7 to −1.6; *P* = 0.011). Switching to isocaloric feeding reduced weight gain during weeks 6–12 (HF/IHF vs HF/HF, −6.4 g, −9.8 to −3.1; *P* < 0.001). IHF/IHF mice had lower glucose (*P* = 0.031) and cholesterol (*P* = 0.005) than HF/HF mice. However, visceral fat relative to body weight remained higher than in LF/LF in IHF/IHF (+1.5 percentage points, 0.04 to 2.9; *P* = 0.042) and HF/IHF (+1.8, 0.4 to 3.3; *P* = 0.008). Adipose *Tnf* expression remained elevated in HF/IHF (log_2_ difference 2.2, 0.2 to 4.2; *P* = 0.025), and fecal phylogenetic diversity was lower in IHF/IHF than in LF/LF (Faith’s PD −3.1, −5.9 to −0.4; *P* = 0.024). Ileal barrier genes and plasma lipopolysaccharide-binding protein were unchanged.

**Conclusions:** Limiting HFD energy intake normalized body weight, glycemia, and plasma lipids but not relative adiposity, adipose *Tnf* expression, or the loss of gut microbial diversity. Diet composition affects these outcomes independently of energy intake, so normal body weight does not indicate their reversal.

## Introduction

Obesity develops when energy intake chronically exceeds expenditure, but diet composition may determine where surplus energy is stored and how tissues respond. In rodents, high-fat diets (HFDs) are the most widely used means of inducing obesity [1]. Mice fed an HFD ad libitum consume more energy than mice fed a low-fat (LF) diet, largely because of the higher energy density and palatability of the HFD, and dietary fat has been identified as a principal driver of energy intake and adiposity in mice [2, 3]. HFD feeding therefore changes both how much and what is eaten, and the phenotypes usually attributed to it, including weight gain, adiposity, adipose tissue inflammation, and altered gut microbiota, could arise from excess energy, from diet composition, or from both. In humans, removing the same number of calories from fat or from carbohydrate produces different losses of body fat [4], indicating that composition matters even when energy is controlled.

The metabolic burden of obesity is not fully captured by body weight. Excess body fat and increased cardiometabolic risk can occur at a normal body mass index [5], and adverse outcomes depend on how adipose tissue expands. Hypertrophic expansion of adipocytes is accompanied by local hypoxia, adipocyte death, and recruitment of macrophages that produce inflammatory cytokines such as tumor necrosis factor-α (TNF-α) [6–8], which impairs insulin action [9, 10]. Saturated fatty acids, abundant in lard-based HFDs, can also activate Toll-like receptor 4 (TLR4) signaling in adipocytes and macrophages directly [11], raising the possibility that adipose inflammation is driven partly by dietary fat independently of energy surplus.

HFD feeding also rapidly remodels the gut microbiota and reduces its diversity [12, 13]. These changes, together with increased intestinal permeability and translocation of bacterial lipopolysaccharide, have been proposed to link diet to adipose inflammation [14, 15]. However, several HFD-induced microbial changes occur independently of obesity [13, 16], and many can be explained by the low fermentable fiber content of purified HFDs rather than by fat itself [17, 18]. Conversely, caloric restriction itself reshapes the microbiota [19–21]. Whether HFD-induced microbial changes persist when energy intake is normalized remains unresolved.

Pair-feeding an HFD to the energy intake of LF-fed mice dissociates diet composition from energy intake [22, 23]. Such studies have mostly matched intake from the onset of HFD feeding, which tests prevention, and have focused on body weight and the liver. Whether matching energy intake after obesity has developed reverses established changes in adipose tissue and the microbiota, a question more relevant to dietary treatment, has received less attention. It is important because obesity-associated immune and epigenetic changes in adipose tissue can persist after weight loss [24–26], and post-dieting microbial alterations can promote weight regain [27].

Here, we restricted HFD intake to the energy intake of LF-fed mice either from the start of feeding (prevention) or after 6 weeks of ad libitum HFD feeding (treatment) and examined body weight, adiposity, circulating metabolic markers, adipose and ileal gene expression, and the fecal microbiota in male C57BL/6 mice. We hypothesized that outcomes driven by excess energy intake would be normalized by isocaloric restriction, whereas outcomes driven by diet composition would persist despite normal body weight.

## Materials and methods

### Animals and ethics

Male C57BL/6 mice (6 weeks old; Samtako Co., Osan, Republic of Korea) were housed individually at 23 ± 1 °C and 50 ± 5% relative humidity under a 12-h light/dark cycle with free access to water. After 3 weeks of acclimatization on the LF diet, the experimental diets were introduced (week 0). All procedures were approved by the Institutional Animal Care and Use Committee of Sun Moon University (approval no. SM-2022-08-01; 24 November 2022) and are reported in accordance with the ARRIVE guidelines [28].

### Diets

The LF (10% of energy from fat) and HF (45% of energy from fat) diets were formulated according to the published compositions of the D12450J and D12451 purified rodent diets, respectively (Supplementary Table 1). Both provided 20% of energy from protein. Besides fat (lard and soybean oil), they differed in carbohydrate source: the HF diet contained more sucrose (17.0% vs 6.8% of energy) and less corn starch (7.2% vs 49.9% of energy). Energy densities calculated from the ingredient composition were 3.85 (LF) and 4.73 kcal/g (HF).

### Experimental design

Thirty-two mice were randomly allocated within body-weight strata to four groups (n = 8) and fed for 12 weeks: LF diet ad libitum (LF/LF); HF diet ad libitum (HF/HF); HF diet ad libitum for 6 weeks, then isocaloric HF feeding (HF/IHF; treatment model); or isocaloric HF feeding throughout (IHF/IHF; prevention model) (Fig. 1A). Body weight was recorded weekly, and food intake was measured daily and converted to energy intake using the energy densities above.

**Fig. 1.**
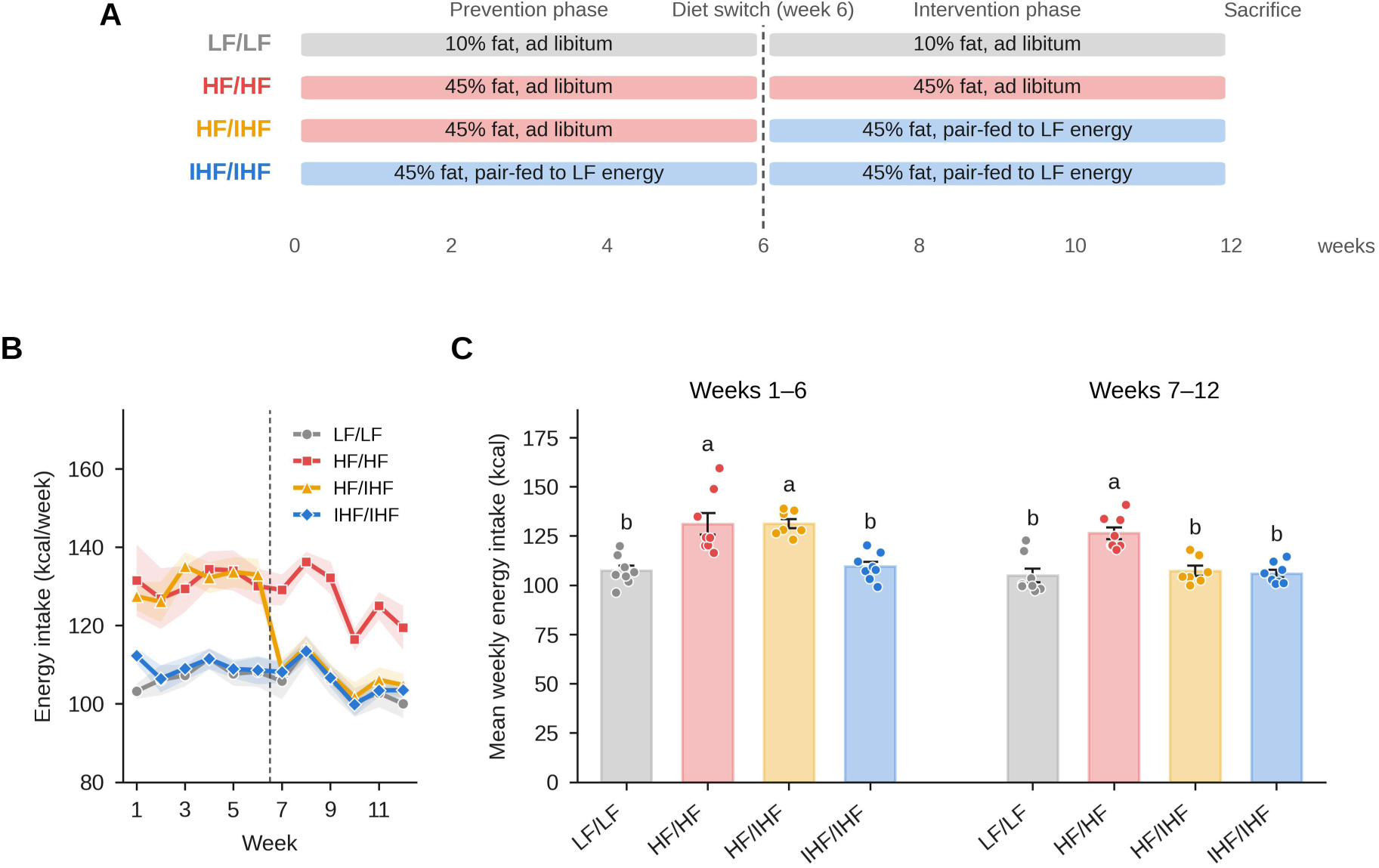
Study design and energy intake. (A) Male C57BL/6 mice were fed a 10% fat diet (LF) or a 45% fat diet (HF) ad libitum, or the HF diet pair-fed to the energy intake of the LF group (IHF), for 6 weeks (prevention phase). After week 6, one HF group was switched to IHF (HF/IHF) and all other groups continued their diets for weeks 7–12 (intervention phase). (B) Weekly energy intake. The dashed line marks the diet switch. (C) Mean weekly energy intake during weeks 1–6 and weeks 7–12. Data are mean ± SEM with individual mice shown (n = 7–8). Bars not sharing a letter differ significantly (one-way ANOVA with Tukey’s test, *P* < 0.05).

### Isocaloric pair-feeding

Each IHF/IHF mouse was yoked to an LF/LF mouse of the same body-weight rank and received daily an amount of HF diet equal to the previous day’s intake of its partner multiplied by the ratio of the LF to HF energy density (≈ 0.8), so that both consumed the same energy. From week 7, each HF/IHF mouse received the same daily ration as one IHF/IHF mouse and was thus matched to the same LF/LF partner. Food intake was calculated as the amount offered minus the amount remaining in the feeder; spillage was not collected separately. Intake data from one HF/IHF mouse were excluded because its recorded intake during weeks 1–3 was implausibly high (up to 84 g/week), indicating food wastage; its other data were retained. No animals were excluded from the analyses.

### CCK-induced satiety test

During week 11 (15–18 February 2023), the anorexigenic response to cholecystokinin (CCK) was assessed. After a 6-h fast, mice received an intraperitoneal injection of saline or sulfated CCK-8 (3 µg/kg body weight; Bachem, Bubendorf, Switzerland), and pre-weighed food was provided immediately. Food intake was measured 20, 40, and 60 min after injection and converted to energy intake, with 40 min as the primary endpoint. Each mouse received both treatments in a crossover design with a washout of at least 48 h, and injection order was counterbalanced across animals. Energy intake at 40 min was analyzed with a linear mixed model with group, treatment, and their interaction as fixed effects and mouse as a random effect; within-group comparisons used paired *t* tests.

### Sample collection

At the end of week 12 (22–23 February 2023), food was removed at approximately 07:00, and mice were killed from 13:00 (≈ 6-h fast) in an order that interleaved the groups. Blood glucose was measured in tail-vein blood with a glucometer. Mice were then euthanized by CO_2_ inhalation, and blood was collected by cardiac puncture; plasma was separated by centrifugation (1,000 × g, 10 min, 4 °C). The liver, epididymal and retroperitoneal fat pads, and cecum were excised and weighed, and portions of fat together with the ileum were frozen on dry ice and stored at −80 °C.

### Plasma analyses

Plasma alanine aminotransferase (ALT), aspartate aminotransferase (AST), total cholesterol, and triglycerides were measured by Dooyeol Biotech (Seoul, Republic of Korea) with an automated analyzer (Hitachi 7180; Hitachi, Tokyo, Japan). Five hemolyzed samples were retained and are marked in Fig. 4. Plasma lipopolysaccharide-binding protein (LBP) was measured in 1:20-diluted samples with a mouse LBP ELISA kit (Elabscience, Houston, TX, USA), and absorbance was read at 450 nm (SPECTROstar Nano; BMG Labtech, Ortenberg, Germany).

### Adipose histology

Epididymal fat was fixed in 10% neutral buffered formalin for 24 h, embedded in paraffin, sectioned at 5 µm, and stained with hematoxylin and eosin. The diameters of approximately 20 adipocytes per mouse were measured in two images (10×; Ki3000 microscope, Korea Lab Tech, Seongnam, Republic of Korea) with ImageJ (NIH, Bethesda, MD, USA).

### Quantitative real-time PCR

Total RNA was extracted from epididymal fat and ileum with TRIzol reagent (Invitrogen, Carlsbad, CA, USA), and cDNA was synthesized with TOPscript RT DryMIX (dT18 plus; Enzynomics, Daejeon, Republic of Korea). Quantitative PCR was performed with TOPreal qPCR 2× PreMIX SYBR Green (Enzynomics) on a StepOnePlus Real-Time PCR System (Thermo Fisher Scientific, Waltham, MA, USA) using the primers listed in Supplementary Table 3. Expression was normalized to *Gapdh* and calculated by the 2^−ΔΔCt^ method relative to the LF/LF mean [29].

### Fecal microbiota analysis

Feces collected on 21 February 2023 (end of week 12, 1 day before the first mice were killed) were analyzed (n = 4 per group). DNA was extracted with the ZR Fecal DNA MiniPrep kit (Zymo Research, Irvine, CA, USA), and the V3–V4 region of the 16S rRNA gene was amplified with primers Bakt_341F and Bakt_805R and sequenced on an Illumina MiSeq platform (2 × 301 bp; Macrogen, Seoul, Republic of Korea). Adapter and primer sequences were removed with Cutadapt v3.2 [30], and amplicon sequence variants (ASVs) were inferred with DADA2 v1.18.0 [31] after quality filtering, denoising, merging, and chimera removal. Taxonomy was assigned against the NCBI 16S rRNA database. ASVs were aligned with MAFFT v7.475, and a phylogenetic tree was built with FastTreeMP v2.1.10 [32, 33]. After rarefaction to 23,800 reads per sample (the smallest library), alpha diversity (observed ASVs, Chao1, Shannon index, and Faith’s phylogenetic diversity) and beta diversity (Bray–Curtis and unweighted and weighted UniFrac distances) were computed with QIIME v1.9.0 [34, 35].

### Statistical analysis

Data are presented as mean ± SEM, with the individual mouse as the experimental unit. Groups were compared by one-way ANOVA followed by Tukey’s multiple comparisons test, with results shown as compact letter displays. Gene expression data were analyzed after log_2_ transformation. Normality of residuals and homogeneity of variance were assessed with the Shapiro–Wilk and Levene tests; where these were violated (energy intake during weeks 7–12, adipose *Tnf*, observed ASVs, and Shannon index), the Kruskal–Wallis test was applied as a sensitivity analysis. One-way ANOVA and Tukey’s tests were performed in GraphPad Prism (version 10.3.1; GraphPad Software, Boston, MA, USA). Body-weight trajectories were analyzed with a linear mixed model with group × week as fixed effects and a random intercept and slope for each mouse. Beta diversity was tested by PERMANOVA [36] with 9,999 permutations, followed by pairwise PERMANOVA. Genera with a mean relative abundance ≥ 0.3% were compared by the Kruskal–Wallis test with Benjamini–Hochberg false discovery rate correction [37], and *Akkermansia* by ANOVA on log-transformed abundance. Genus–phenotype associations were explored by Spearman correlation without multiplicity correction. These analyses were performed in Python 3.11 with SciPy 1.17, statsmodels 0.15, and scikit-bio 0.7. Two-sided *P* < 0.05 was considered significant.

## Results

### Isocaloric pair-feeding matched the energy intake of HF- and LF-fed mice

During weeks 1–6, mice fed the HF diet ad libitum consumed approximately 22% more energy than LF/LF mice (131.0 ± 5.5 and 131.2 ± 2.4 vs 107.3 ± 2.6 kcal/week; both *P* < 0.001), whereas IHF/IHF mice consumed 109.4 ± 2.4 kcal/week (*P* = 0.97 vs LF/LF; Fig. 1B, C). After the diet switch, HF/IHF mice consumed 107.2 ± 2.6 kcal/week, similar to LF/LF (104.9 ± 3.4) and IHF/IHF (105.9 ± 1.9) mice, whereas HF/HF mice continued to overconsume (126.4 ± 3.0 kcal/week; *P* < 0.001 vs each other group; Fig. 1C). In a satiety test at week 11, CCK-8 reduced 40-min energy intake overall (*P* = 0.029), with no evidence that the response differed among groups (group × treatment interaction *P* = 0.29; Supplementary Table 4).

### Isocaloric restriction prevented weight gain and halted further gain in mice with diet-induced obesity

Initial body weight did not differ among groups (26.4–26.7 g; *P* = 0.99). The rate of weight gain was lower in LF/LF, HF/IHF, and IHF/IHF mice than in HF/HF mice (group × week interaction, −0.58, −0.50, and −0.59 g/week, respectively; all *P* < 0.001; Fig. 2A). In the prevention model, IHF/IHF mice gained as much weight as LF/LF mice during weeks 0–6 (7.4 ± 1.0 vs 7.9 ± 0.8 g), about half the gain of mice fed the HF diet ad libitum (13.1–14.7 g; Fig. 2C), and at week 12 the two groups had the same body weight (39.0 ± 1.5 vs 38.9 ± 1.0 g; *P* > 0.99), both lower than that of HF/HF mice (47.2 ± 2.2 g; *P* = 0.011 and 0.009; Fig. 2B). In the treatment model, isocaloric restriction almost completely stopped further gain during weeks 6–12 (HF/IHF, 1.1 ± 1.3 g vs HF/HF, 7.5 ± 0.7 g; *P* < 0.001), to below the gain of LF/LF mice (4.5 ± 0.5 g; *P* = 0.048; Fig. 2C); the final body weight of HF/IHF mice (42.5 ± 1.9 g) was intermediate. Feed efficiency was higher in HF/HF mice (1.33 ± 0.07 g/100 kcal) than in LF/LF and IHF/IHF mice (both 0.97 g/100 kcal; *P* = 0.028 and 0.029; Supplementary Table 2).

**Fig. 2.**
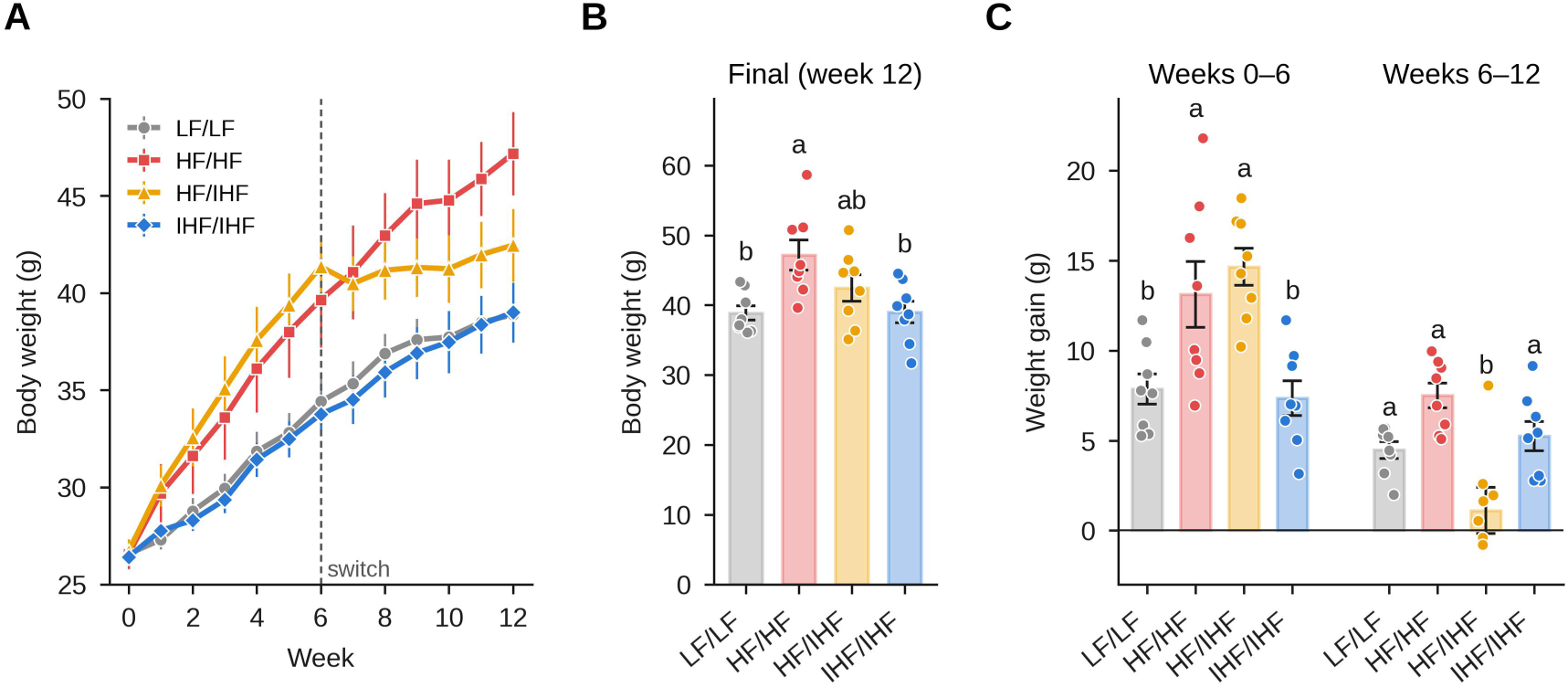
Isocaloric restriction prevents HFD-induced weight gain and halts further gain in mice with diet-induced obesity. (A) Body weight over 12 weeks. (B) Final body weight. (C) Weight gain during weeks 0–6 and weeks 6–12. Data are mean ± SEM (n = 7–8). Bars not sharing a letter differ significantly (one-way ANOVA with Tukey’s test, *P* < 0.05).

### Isocaloric HF feeding increased relative adiposity despite normal body weight

Visceral fat (retroperitoneal plus epididymal) as a proportion of body weight was higher in HF/HF (7.1 ± 0.4%), HF/IHF (7.4 ± 0.3%), and IHF/IHF (7.0 ± 0.4%) mice than in LF/LF mice (5.5 ± 0.3%; *P* = 0.024, 0.008, and 0.042, respectively), with no difference among the HF-fed groups (Fig. 3C). Absolute visceral fat mass was greater in HF/HF and HF/IHF than in LF/LF mice (*P* = 0.003 and 0.022) and intermediate in IHF/IHF mice (Supplementary Table 2). Epididymal adipocytes were larger in HF/IHF than in LF/LF mice (79.4 ± 2.8 vs 66.4 ± 3.7 µm; *P* = 0.043), with a similar trend in HF/HF mice (77.8 ± 3.3 µm; *P* = 0.091) and an intermediate diameter in IHF/IHF mice (70.7 ± 3.3 µm; Fig. 3A, B). Relative liver weight was lowest in IHF/IHF mice (3.03 ± 0.12% vs 4.13 ± 0.26% in LF/LF and 3.97 ± 0.29% in HF/HF; *P* = 0.016 and 0.045; Supplementary Table 2), and relative cecum weight was lower in both restricted groups than in LF/LF mice (0.43 ± 0.05% and 0.42 ± 0.02% vs 0.64 ± 0.03%; *P* = 0.002 and 0.001; Supplementary Table 2).

**Fig. 3.**
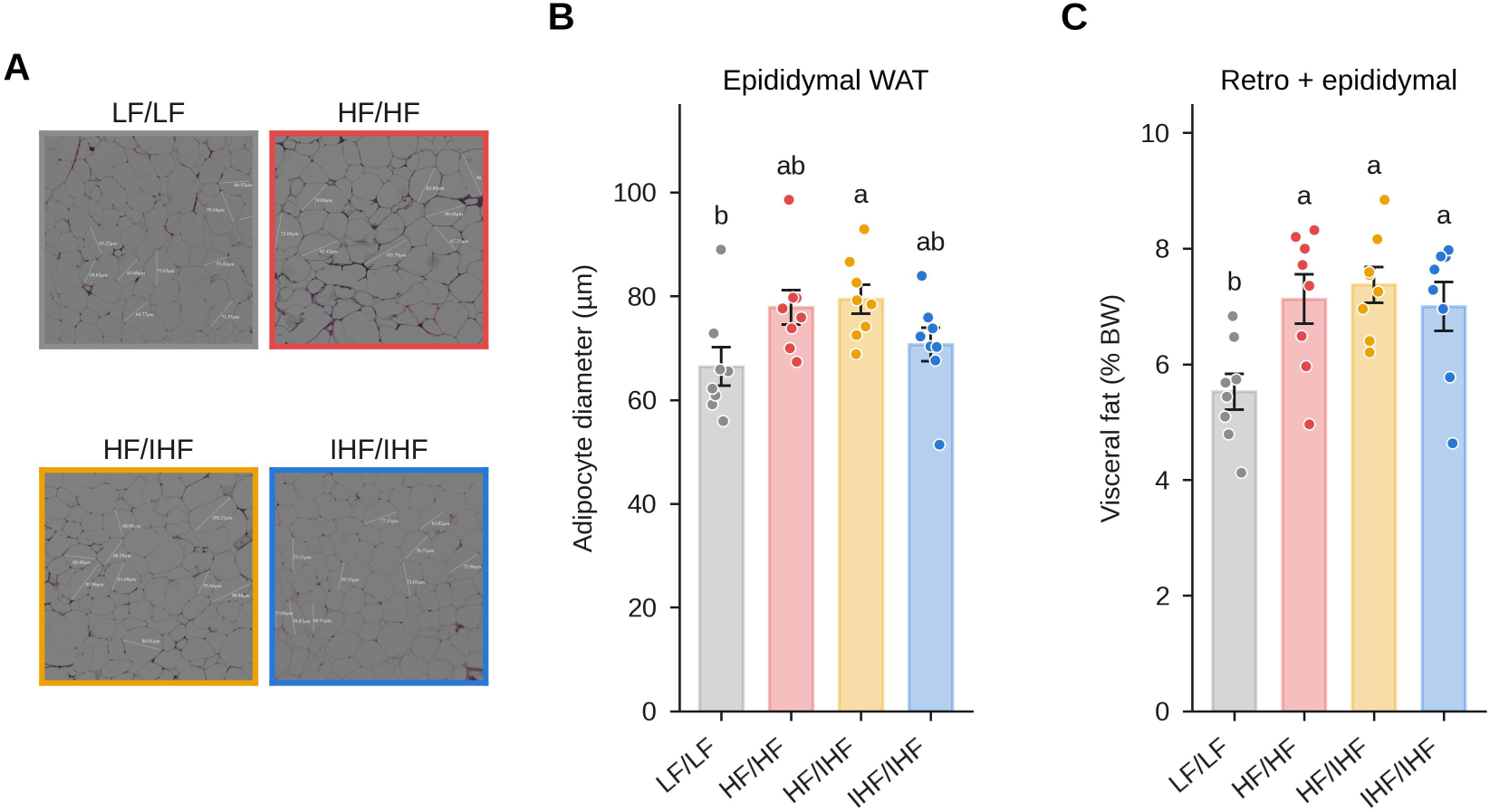
Isocaloric HF feeding increases relative adiposity despite normal body weight. (A) Representative H&E-stained sections of epididymal white adipose tissue (WAT). (B) Mean adipocyte diameter (20 adipocytes per mouse). (C) Combined retroperitoneal and epididymal fat mass relative to body weight. Data are mean ± SEM (n = 8). Bars not sharing a letter differ significantly (one-way ANOVA with Tukey’s test, *P* < 0.05).

### Isocaloric restriction improved glycemia and plasma lipids

Blood glucose was highest in HF/HF mice (194 ± 13 mg/dL) and was lower in IHF/IHF mice (151 ± 9 mg/dL; *P* = 0.031), with a similar trend in HF/IHF mice (157 ± 12 mg/dL; *P* = 0.076; Fig. 4A). Plasma total cholesterol was higher in HF/HF than in LF/LF mice (234 ± 16 vs 178 ± 11 mg/dL; *P* = 0.024) and was normalized in IHF/IHF mice (167 ± 13 mg/dL; *P* = 0.005 vs HF/HF), whereas HF/IHF mice were intermediate (212 ± 11 mg/dL; Fig. 4B). Plasma triglycerides were lower in HF/IHF (27 ± 4 mg/dL) and IHF/IHF (22 ± 3 mg/dL) than in HF/HF mice (52 ± 9 mg/dL; *P* = 0.045 and 0.009; Fig. 4C), and ALT and AST did not differ among groups (Supplementary Table 2).

**Fig. 4.**
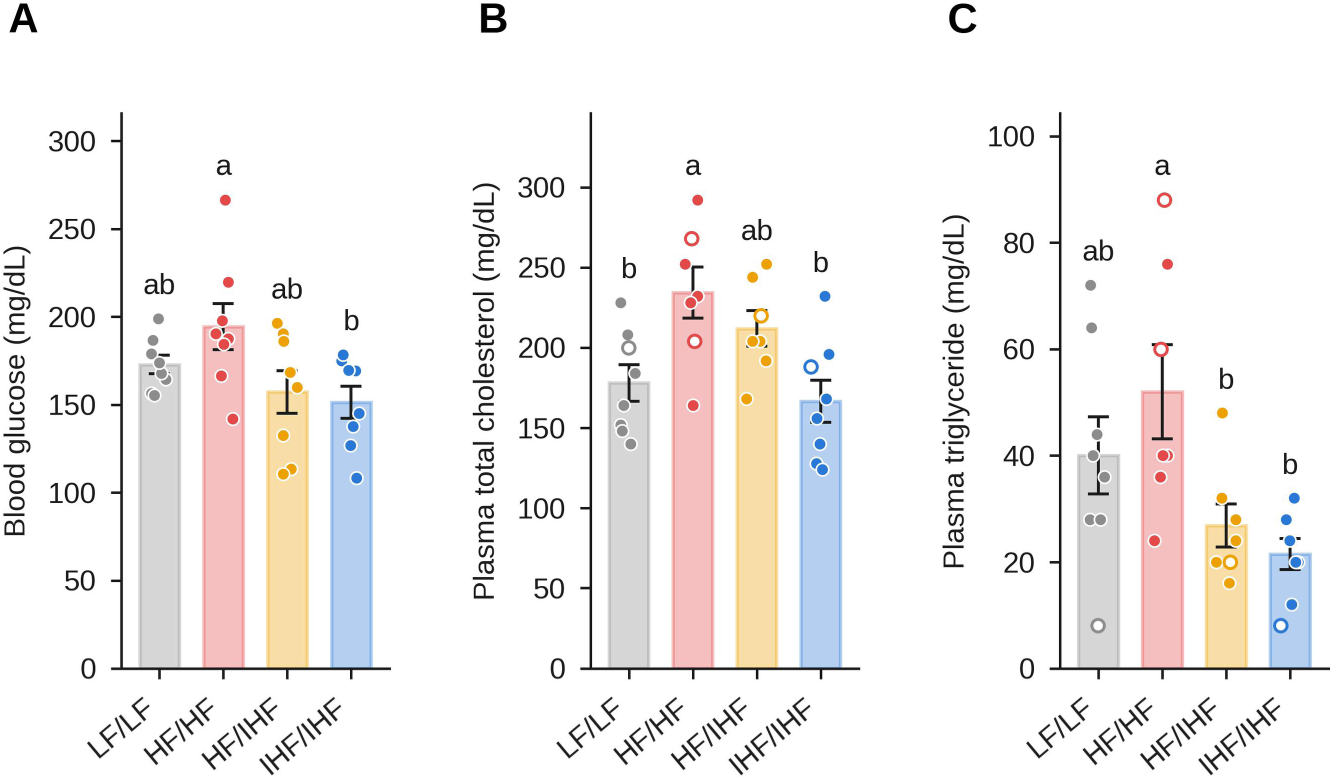
Isocaloric restriction improves glycemia and plasma lipids. (A) Blood glucose at sacrifice. (B) Plasma total cholesterol. (C) Plasma triglycerides. Open circles indicate hemolyzed samples. Data are mean ± SEM (n = 7–8). Bars not sharing a letter differ significantly (one-way ANOVA with Tukey’s test, *P* < 0.05).

### Adipose Tnf expression remained elevated after isocaloric restriction in mice with diet-induced obesity

In epididymal adipose tissue, *Tnf* mRNA was increased 6.4-fold in HF/HF and 8.0-fold in HF/IHF mice relative to LF/LF mice (*P* = 0.009 and 0.025); IHF/IHF mice showed an intermediate 4.8-fold increase (*P* = 0.065), and the three HF-fed groups did not differ from one another (Fig. 5A). *Il10* was increased approximately 4-fold in HF/HF and HF/IHF mice (both *P* = 0.010) but not in IHF/IHF mice (1.5-fold; *P* = 0.89). *Il6*, *Ccl2*, *Lcn2*, *Tlr4*, *Cd14*, and *Pparg* tended to be higher in HF-fed mice, but these differences were not significant (ANOVA *P* = 0.066–0.26; Fig. 5). In the ileum, expression of tight-junction (*Ocln*, *Tjp1*, *Cldn1*), mucus and antimicrobial (*Muc2*, *Defb2*), and inflammatory (*Tnf*, *Tlr4*, *Il1b*) genes did not differ among groups (all ANOVA *P* ≥ 0.055), and plasma LBP was unchanged (*P* = 0.63; Supplementary Table 2).

**Fig. 5.**
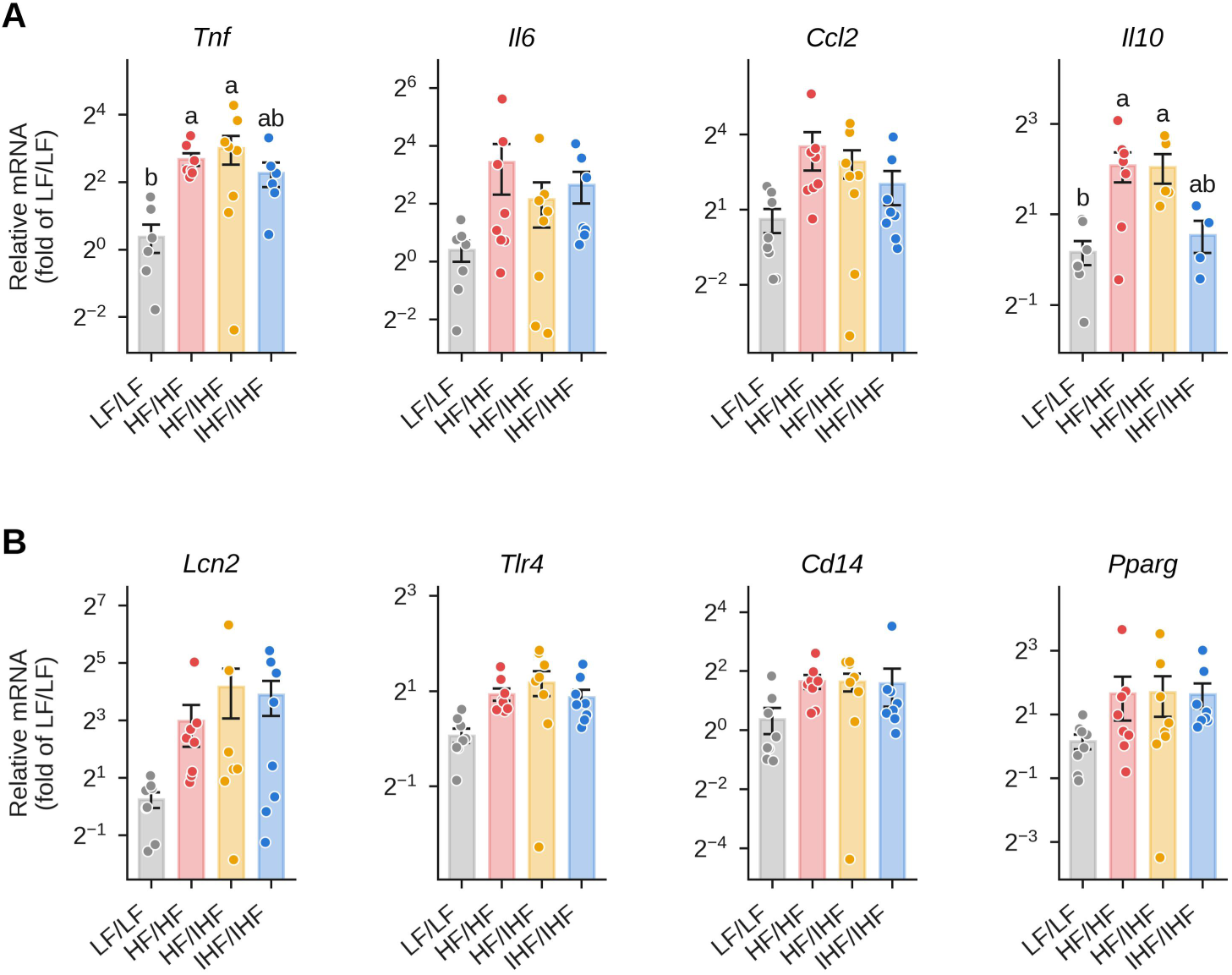
Adipose *Tnf* expression is not normalized by isocaloric restriction. mRNA expression in epididymal WAT of (A) inflammatory cytokines and chemokines (*Tnf*, *Il6*, *Ccl2*, *Il10*) and (B) lipopolysaccharide-sensing and adipogenic genes (*Lcn2*, *Tlr4*, *Cd14*, *Pparg*), normalized to *Gapdh* and expressed as fold change relative to LF/LF (2^−ΔΔCt^). Data are mean ± SEM on a log2 scale (n = 4–8). Statistics were performed on log2-transformed values. Bars not sharing a letter differ significantly (one-way ANOVA with Tukey’s test, *P* < 0.05); genes without letters did not differ by ANOVA.

### Isocaloric restriction did not restore gut microbial diversity

Sequencing yielded 23,800–44,878 high-quality reads per sample. Observed ASVs were lower in HF/HF (237 ± 16) and HF/IHF (203 ± 3) than in LF/LF mice (306 ± 10; *P* = 0.002 and < 0.001; Fig. 6A). IHF/IHF mice retained more ASVs (271 ± 7) than HF/IHF mice ( *P* = 0.002), but their Chao1 richness remained below that of LF/LF mice (272 ± 8 vs 314 ± 11; *P* = 0.043; Supplementary Fig. 1A). Faith’s phylogenetic diversity was reduced to a similar extent in all HF-fed groups (17.9–19.9 vs 23.0; *P* ≤ 0.024; Fig. 6A), whereas the Shannon index did not differ by ANOVA (*P* = 0.17; Kruskal–Wallis *P* = 0.027, reflecting slightly higher values in IHF/IHF mice; Supplementary Fig. 1A).

**Fig. 6.**
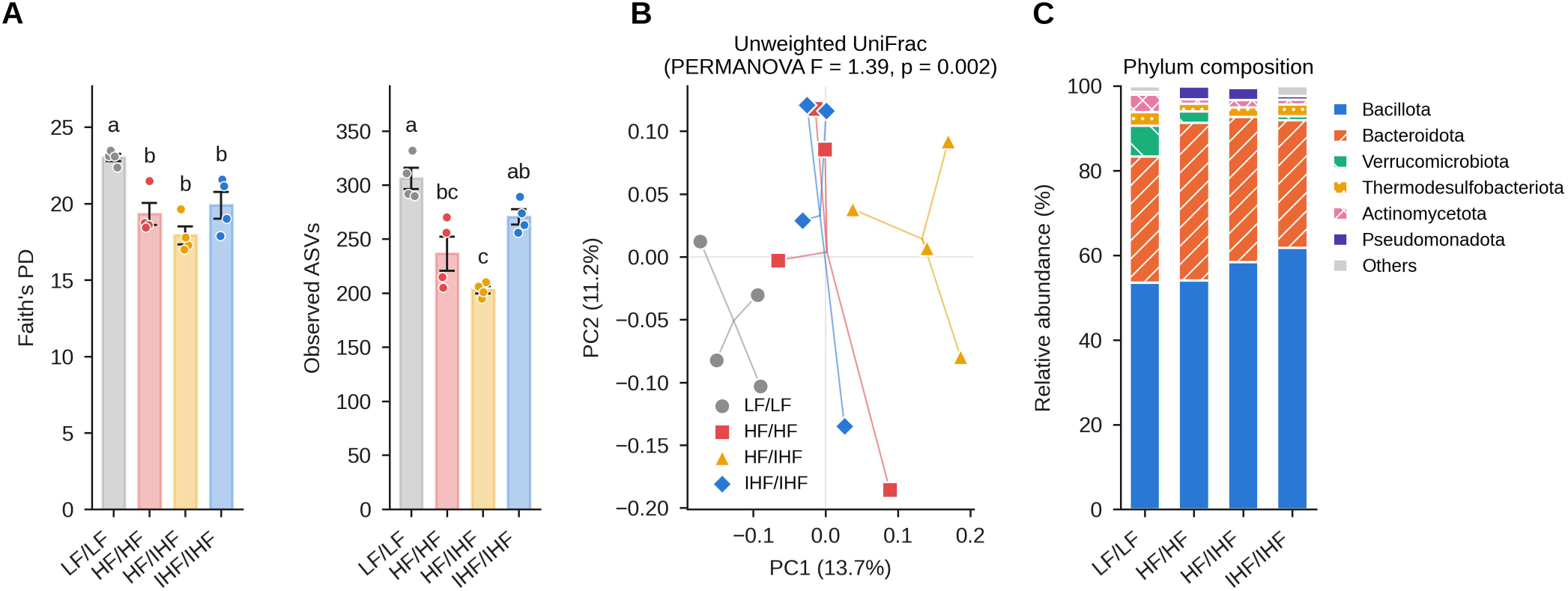
Gut microbial richness and phylogenetic diversity are not restored by isocaloric restriction. Fecal microbiota were analyzed by 16S rRNA gene (V3–V4) sequencing (n = 4 per group). (A) Faith’s phylogenetic diversity and observed ASVs (mean ± SEM with individual mice shown); bars not sharing a letter differ significantly (one-way ANOVA with Tukey’s test, *P* < 0.05). (B) Principal coordinate analysis based on unweighted UniFrac distances. Lines connect samples to group centroids. PERMANOVA, 9,999 permutations. (C) Phylum-level composition (group means; six most abundant phyla).

Community composition differed among groups by unweighted UniFrac (PERMANOVA F = 1.39, *P* = 0.002) and Bray–Curtis distances (F = 1.38, *P* = 0.04) but not by weighted UniFrac distances (*P* = 0.62). Each HF-fed group separated from LF/LF mice on unweighted UniFrac (pairwise *P* = 0.028, the minimum attainable with n = 4), whereas HF/HF and IHF/IHF mice did not differ (*P* = 0.70; Fig. 6B). Phylum-level composition did not differ significantly (Fig. 6C).

*Acutalibacter* (Kruskal–Wallis *P* = 0.027) and *Desulfovibrio* (*P* = 0.060) were depleted in all HF-fed groups, whereas *Lactococcus*, *Lacrimispora*, and *Sangeribacter* showed the opposite pattern (*P* = 0.028–0.030; Supplementary Fig. 1B); none remained significant after false discovery rate correction (*q* ≥ 0.29). *Akkermansia* was nearly absent in HF/IHF mice (0.04% vs 7.2% in LF/LF; *P* = 0.051). Across the 16 sequenced mice, no genus correlated with body weight, whereas *Acutalibacter* correlated negatively with adipocyte size (ρ = −0.65, *P* = 0.006) and adipose *Tnf* expression (ρ = −0.78, *P* = 0.002), *Sangeribacter* positively with adipocyte size (ρ = 0.74, *P* = 0.001), and *Lactococcus* positively with visceral fat (ρ = 0.63, *P* = 0.008; Supplementary Fig. 1C).

## Discussion

Using yoked pair-feeding, we separated the energy content of an HFD from its composition in prevention and treatment settings. Restricting HFD intake to the energy intake of LF-fed mice normalized body weight, glycemia, and plasma lipids but not the proportion of body weight stored as visceral fat, adipose *Tnf* expression in mice with established obesity, or fecal microbial phylogenetic diversity. Body weight and circulating metabolic markers therefore followed energy intake, whereas adiposity, adipose inflammation, and microbial diversity remained associated with diet composition despite energy matching.

IHF/IHF mice gained the same weight as LF/LF mice with identical feed efficiency, so the HFD did not promote weight gain per kcal, and the excess weight of HF/HF mice reflected their approximately 20% greater energy intake. Dietary fat thus promotes weight gain in mice chiefly by increasing intake [3]. Pair-feeding studies that reported greater weight gain on an HFD at matched intake used diets providing 60% of energy from fat [22, 23], suggesting that any effect of fat on energy efficiency is dose dependent. In the treatment model, restriction halted further gain, and HF/IHF mice even gained less than LF/LF mice during weeks 7–12, but it did not reverse the weight already gained within 6 weeks.

At matched energy intake and body weight, IHF/IHF mice carried the same visceral fat proportion as HF/HF mice. Because their absolute fat mass was intermediate and their liver weight was the lowest of all groups, this probably reflects both preferential fat storage and a smaller lean or organ mass. Dietary fat is stored at a lower energetic cost than carbohydrate, which must first be converted to lipid [38], and in humans the macronutrient removed during energy restriction determines how much body fat is lost [4]. A normal body weight on an HFD therefore did not signify normal adiposity, a dissociation that parallels normal-weight obesity in humans [5]. In the treatment model, HF/IHF mice retained both a high fat proportion and the largest adipocytes, indicating that halting weight gain did not reverse adipose expansion within 6 weeks.

Glycemia and lipids, by contrast, followed energy intake. Glucose and cholesterol were lower in IHF/IHF than in HF/HF mice, and triglycerides were lower in both restricted groups, consistent with the metabolic benefits of calorie restriction on an HFD [39]. The prolonged daily fast imposed by pair-feeding may have contributed, and because insulin was not assessed, whether insulin sensitivity was restored remains unknown.

Adipose *Tnf* expression in HF/IHF mice remained as high as in HF/HF mice despite 6 weeks of energy-matched feeding. Hypertrophic adipocytes promote hypoxia, adipocyte death, and macrophage recruitment [6–8], and HF/IHF mice retained the largest adipocytes, which may have sustained this inflammatory milieu. The finding also accords with evidence that obesity-associated immune, transcriptional, and epigenetic features of adipose tissue persist after weight loss [24, 25]; in mice switched from an HFD to chow, adipose inflammation and fibrosis likewise persisted after body weight had normalized [26]. Such obesogenic memory may predispose to relapse. In IHF/IHF mice, *Tnf* expression and adipocyte size were intermediate. Saturated fatty acids from lard can activate TLR4 signaling independently of obesity [11], which may account for a partial composition-driven effect, although adipose *Tlr4* expression did not differ significantly. *Il10* rose only in the groups fed the HFD ad libitum, possibly reflecting the macrophage accumulation that accompanies adipose expansion [10, 40].

These changes occurred without evidence of gut barrier disruption: ileal tight-junction, mucus, and antimicrobial gene expression and plasma LBP were unaltered. HFD-induced metabolic impairment can develop without loss of barrier integrity [41], and our data suggest that metabolic endotoxemia [14] was not a major driver of adipose *Tnf* expression in this model, although the colon, in vivo permeability, and circulating endotoxin were not assessed. In an exploratory test, exogenous CCK reduced short-term intake similarly across groups, suggesting that this gut–brain satiety signal was not grossly impaired, although the low post-fast intake of LF/LF mice limited the assay.

Fecal phylogenetic diversity was reduced in all HF-fed groups irrespective of energy intake, and the HF/HF and IHF/IHF communities were indistinguishable, indicating that microbial diversity and community structure remained associated with diet composition despite energy matching [13, 16]. Differences lay mainly in low-abundance lineages (unweighted but not weighted UniFrac). The relevant dietary component may not be fat itself: the LF diet supplied approximately seven times more energy from corn starch, part of which reaches the hindgut as fermentable substrate [42], and many microbial effects attributed to HFDs are reproduced by removing fermentable fiber [17, 18]. Loss of fermentable carbohydrate reduces diversity [43], favors mucus degradation [44], and impairs IL-22-dependent colonic health [45]. Caloric restriction itself also reshapes the microbiota [19–21]; together with prior HFD exposure, this may explain why HF/IHF mice had the lowest richness and almost no *Akkermansia*, a mucus-associated genus that protects against diet-induced metabolic disorders [46]. Taxa altered in all HF-fed groups correlated with adiposity and adipose *Tnf* expression but not with body weight, although these exploratory associations require confirmation. Because persistent microbial changes can accelerate weight regain after dieting [27], failure to restore the microbiota may contribute to relapse after energy restriction alone.

Strengths of this study include daily yoked pair-feeding, which kept energy intake within approximately 2% of that of LF-fed mice in both settings, and the inclusion of prevention and treatment arms in one experiment. Several limitations should be considered. First, the LF diet was not matched to the HF diet in sucrose and starch; a sucrose-matched control (D12450H) would have isolated the effect of fat, so our findings apply to diet composition as a whole. Second, pair-fed mice consume their ration within a few hours, imposing a longer daily fast [47]; because time-restricted feeding alone improves metabolism on an HFD [48], some effects may reflect altered feeding patterns. Third, the microbiota was analyzed in four mice per group at one time point. Fourth, only male mice were studied, and body composition, energy expenditure, and insulin sensitivity were not measured. Finally, 6 weeks may be too short to reverse adipose remodeling.

In conclusion, isocaloric restriction of an HFD normalized body weight, glycemia, and plasma lipids but not the visceral fat proportion, adipose *Tnf* expression after established obesity, or microbial diversity, which depended on diet composition independently of energy intake. These findings indicate that normalization of body weight does not necessarily imply normalization of adiposity or of diet-associated adipose and microbial phenotypes. Whether this dissociation extends to humans requires direct investigation. Sucrose-matched and fiber-supplemented diets, longer restriction, and female mice should be examined next.

## Data availability

Raw 16S rRNA amplicon sequencing data generated in this study have been deposited in the NCBI Sequence Read Archive (SRA) under BioProject accession number PRJNA1535893 and are publicly available at https://www.ncbi.nlm.nih.gov/bioproject/PRJNA1535893. All other data supporting the findings of this study are available from the corresponding author upon reasonable request.

## Author contributions

Sungjin Eun: Data curation, Formal analysis, Investigation, Methodology, Visualization, Writing – original draft. Eunji Jeong: Data curation, Formal analysis, Investigation, Methodology, Writing – review & editing. Seoyeon Chae: Data curation, Formal analysis, Investigation, Methodology, Writing – review & editing. Sunhye Lee: Conceptualization, Formal analysis, Funding acquisition, Investigation, Methodology, Project administration, Resources, Supervision, Validation, Visualization, Writing – original draft, Writing – review & editing.

## Ethics approval

All animal procedures were approved by the Institutional Animal Care and Use Committee of Sun Moon University (approval no. SM-2022-08-01) and conducted in accordance with the ARRIVE guidelines.

