## Supplementary Tables 1-4 and Supplementary Figure 1 for "Isocaloric restriction normalizes body weight but not relative adiposity, adipose inflammation, or gut microbial diversity in high-fat diet-fed mice"

**Summary:** This file contains four supplementary tables (experimental diet composition; growth, tissue weights, plasma markers and ileal gene expression; qPCR primer sequences; and energy intake after CCK-8 injection) and one supplementary figure (additional exploratory fecal microbiota analyses). File format: PDF.

**Supplementary Table 1.** Composition of the experimental diets.

| Component | LF diet | HF diet |
|---|---|---|
| **Ingredient (g/kg diet)** | | |
| Casein | 189.6 | 233.1 |
| L-Cystine | 2.8 | 3.5 |
| Corn starch | 479.8 | 84.8 |
| Maltodextrin 10 | 118.5 | 116.5 |
| Sucrose | 65.2 | 201.4 |
| Cellulose | 47.4 | 58.3 |
| Soybean oil | 23.7 | 29.1 |
| Lard | 19.0 | 206.8 |
| Mineral mix S10026 | 9.5 | 11.7 |
| Dicalcium phosphate | 12.3 | 15.1 |
| Calcium carbonate | 5.2 | 6.4 |
| Potassium citrate, 1 $H_2O$ | 15.6 | 19.2 |
| Vitamin mix V10001 | 9.5 | 11.7 |
| Choline bitartrate | 1.9 | 2.3 |
| **Energy** | | |
| Energy density (kcal/g) | 3.85 | 4.73 |
| Protein (% kcal) | 20.0 | 20.0 |
| Carbohydrate (% kcal) | 70.0 | 35.0 |
| Sucrose (% kcal) | 6.8 | 17.0 |
| Corn starch (% kcal) | 49.9 | 7.2 |
| Fat (% kcal) | 10.0 | 45.0 |

LF, low-fat diet (10% of energy from fat); HF, high-fat diet (45% of energy from fat). Formulations follow the published compositions of the D12450J (LF) and D12451 (HF) purified rodent diets. Ingredient amounts are per kg of diet; dyes are omitted. Energy density was calculated from the ingredient composition. Carbohydrate energy includes the sucrose carrier of the vitamin mix. The two diets differ not only in fat but also in sucrose and corn-starch content.

**Supplementary Table 2.** Tissue weights, plasma markers, and ileal gene expression.

| Variable | LF/LF | HF/HF | HF/IHF | IHF/IHF | *P* |
|---|---|---|---|---|---|
| **Growth** | | | | | |
| Feed efficiency (g/100 kcal) | 0.97 ± 0.06$^{b}$ | 1.33 ± 0.07$^{a}$ | 1.08 ± 0.11$^{ab}$ | 0.97 ± 0.10$^{b}$ | 0.018 |
| **Tissue weights** | | | | | |
| Liver (% body weight) | 4.13 ± 0.26$^{a}$ | 3.97 ± 0.29$^{a}$ | 3.57 ± 0.26$^{ab}$ | 3.03 ± 0.12$^{b}$ | 0.015 |
| Visceral fat (g) | 2.16 ± 0.16$^{b}$ | 3.37 ± 0.28$^{a}$ | 3.11 ± 0.15$^{a}$ | 2.77 ± 0.25$^{ab}$ | 0.003 |
| Cecum (% body weight) | 0.64 ± 0.03$^{a}$ | 0.51 ± 0.05$^{ab}$ | 0.43 ± 0.05$^{b}$ | 0.42 ± 0.02$^{b}$ | 0.001 |
| **Plasma** | | | | | |
| LBP (ng/mL) | 313 ± 23 | 318 ± 22 | 310 ± 28 | 279 ± 19 | 0.63 |
| ALT (U/L) | 65.2 ± 20.2 | 64.2 ± 14.0 | 102.2 ± 21.5 | 65.5 ± 15.9 | 0.41 |
| AST (U/L) | 144.7 ± 27.4 | 121.4 ± 14.7 | 185.6 ± 33.5 | 212.2 ± 73.2 | 0.50 |
| **Ileal mRNA (fold of LF/LF)** | | | | | |
| *Ocln* | 1.24 ± 0.25 | 1.35 ± 0.42 | 1.11 ± 0.41 | 1.12 ± 0.27 | 0.53 |
| *Tjp1* | 1.13 ± 0.19 | 0.94 ± 0.19 | 0.82 ± 0.27 | 0.96 ± 0.19 | 0.37 |
| *Cldn1* | 1.06 ± 0.13 | 1.20 ± 0.14 | 1.23 ± 0.17 | 1.20 ± 0.18 | 0.97 |
| *Muc2* | 1.17 ± 0.23 | 1.02 ± 0.20 | 1.02 ± 0.45 | 0.96 ± 0.21 | 0.71 |
| *Defb2* | 1.08 ± 0.15 | 1.15 ± 0.14 | 1.30 ± 0.18 | 1.26 ± 0.18 | 0.90 |
| *Tnf* | 1.08 ± 0.13 | 1.51 ± 0.19 | 1.62 ± 0.22 | 1.68 ± 0.27 | 0.43 |
| *Il1b* | 1.03 ± 0.09 | 1.78 ± 0.23 | 1.62 ± 0.14 | 1.86 ± 0.25 | 0.055 |
| *Tlr4* | 1.08 ± 0.14 | 1.27 ± 0.16 | 1.33 ± 0.19 | 1.41 ± 0.22 | 0.86 |

Data are mean ± SEM (n = 8 per group; n = 7 for HF/HF and HF/IHF plasma enzymes and for HF/IHF feed efficiency). Feed efficiency is body weight gained per 100 kcal consumed over 12 weeks. Visceral fat is the sum of retroperitoneal and epididymal fat pads. Values not sharing a superscript letter differ significantly (Tukey's test, $P < 0.05$). Ileal mRNA levels were normalized to *Gapdh* and expressed as fold change relative to LF/LF ($2^{-\Delta\Delta Ct}$); statistics were performed on $\log_2$-transformed values. *P* values are from one-way ANOVA. ALT, alanine aminotransferase; AST, aspartate aminotransferase; LBP, lipopolysaccharide-binding protein.

**Supplementary Table 3.** Primer sequences used for quantitative real-time PCR.

| Gene | Tissue | Forward (5′→3′) | Reverse (5′→3′) |
|---|---|---|---|
| *Tnf* | eWAT, ileum | GGGTGTTCATCCATTCTCTAC | CCCAGCATCTTGTGTTTCT |
| *Il6* | eWAT | CCAGAGTCCTTCAGAGAGATA | ACTAGGTTTGCCGAGTAGA |
| *Ccl2* | eWAT | TTAAAAACCTGGATCGGAACCAA | GCATTAGCTTCAGATTTACGGGT |
| *Il10* | eWAT | CAGCCGGGAAGACAATAAC | GCTTGGCAACCCAAGTAA |
| *Lcn2* | eWAT | CAACTGAATGGGTGGTGAG | GGAGTGCTGGCCAAATAA |
| *Tlr4* | eWAT, ileum | ATGGCATGGCTTACACCACC | GAGGCCAATTTTGTCTCCACA |
| *Cd14* | eWAT | AGTGAGTAGGGCTGTTAGG | CCGAGAAAGGCTTCAGTTT |
| *Pparg* | eWAT | GGCCTCCCTGATGAATAAAG | CGGTCTCCACTGAGAATAATG |
| *Il1b* | ileum | GGGCTGGACTGTTTCTAATG | TCAGAGGCAAGGAGGAAA |
| *Ocln* | ileum | TTGAAAGTCCACCTCCTTACAGA | CCGGATAAAAAGAGTACGCTGG |
| *Tjp1* | ileum | GCCGCTAAGAGCACAGCAA | TCCCCACTCTGAAAATGAGGA |
| *Cldn1* | ileum | GCTGGGTTTCATCCTGGCTTCT | CCTGAGCGGTCACGATGTTGTC |
| *Muc2* | ileum | ATGCCCACCTCCTCAAAGAC | GTAGTTTCCGTTGGAACAGTGAA |
| *Defb2* | ileum | CACTCCAGCTGTTGGAAGTTTA | GCTCTGACACAGTACCCTCC |
| *Gapdh* | eWAT, ileum | AGGTCGGTGTGAACGGATTTG | TGTAGACCATGTAGTTGAGGTCA |

eWAT, epididymal white adipose tissue.

**Supplementary Table 4.** Energy intake after intraperitoneal saline or CCK-8 (week 11).

| **Group** | **Time (min)** | **Saline (kcal)** | **CCK-8 (kcal)** | ***P*** |
|---|---|---|---|---|
| LF/LF | 20 | 0.06 ± 0.12 | 0.17 ± 0.12 | 0.582 |
| | 40 | 0.38 ± 0.17 | 0.50 ± 0.28 | 0.750 |
| | 60 | 0.51 ± 0.19 | 0.86 ± 0.30 | 0.286 |
| HF/HF | 20 | 0.63 ± 0.08 | 0.30 ± 0.19 | 0.134 |
| | 40 | 0.95 ± 0.09 | 0.64 ± 0.18 | 0.104 |
| | 60 | 1.21 ± 0.15 | 0.86 ± 0.15 | 0.110 |
| HF/IHF | 20 | 1.83 ± 0.45 | 0.87 ± 0.46 | 0.074 |
| | 40 | 2.10 ± 0.43 | 1.33 ± 0.40 | 0.046 |
| | 60 | 2.71 ± 0.72 | 1.92 ± 0.57 | 0.134 |
| IHF/IHF | 20 | 1.45 ± 0.56 | 0.60 ± 0.36 | 0.157 |
| | 40 | 2.03 ± 0.61 | 1.38 ± 0.55 | 0.271 |
| | 60 | 2.41 ± 0.66 | 1.79 ± 0.56 | 0.294 |

Data are cumulative energy intake (mean ± SEM; n = 8 per group), calculated from food intake and diet energy density. Mice were fasted for 6 h before an intraperitoneal injection of saline or sulfated CCK-8 (3 µg/kg) in a counterbalanced crossover design (washout ≥ 48 h). *P* values are from paired *t* tests within each group; 40 min was the primary endpoint. A linear mixed model of 40-min energy intake (group × treatment; mouse as random effect) showed a main effect of CCK-8 ($P = 0.029$) and no group × treatment interaction ($P = 0.29$). These data are exploratory.

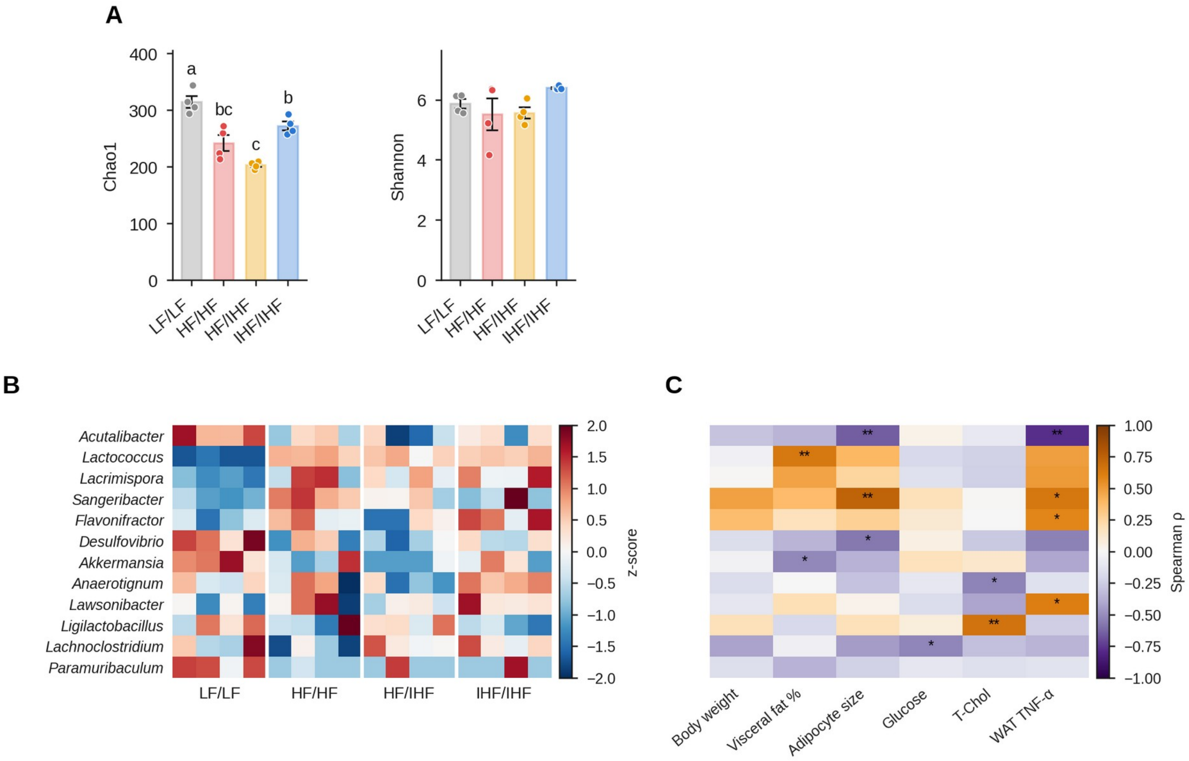


**Supplementary Fig. 1. Additional microbiota analyses (exploratory).** (A) Chao1 richness and Shannon index (mean ± SEM; n = 4 per group). Bars not sharing a letter differ significantly (one-way ANOVA with Tukey's test, $P < 0.05$). (B) Heat map of the 12 genera with the lowest Kruskal–Wallis *P* values (mean relative abundance ≥ 0.3%), shown as row z-scores of log10 abundance; none remained significant after false discovery rate correction ($q \geq 0.29$). (C) Spearman correlations between these genera (rows as in B) and host phenotypes across the 16 sequenced mice. *$P < 0.05$, **$P < 0.01$ (uncorrected).
